# Reconstructing their genomes confirms the historically attested genealogy of the two medieval emperors Otto I (“the Great”) and Heinrich II (“Saint Henry”)

**DOI:** 10.64898/2026.03.18.712637

**Authors:** Harald Ringbauer, Thomas Wozniak, Jörg Feuchter, Göran Runfeldt, Raffaela Angelina Bianco, Ganyu Zhang, Kay Prüfer, Jörg Orschiedt, Annika Simm, Paul Maier, Michael Sager, Veit Dresely, Johannes Krause, Harald Meller, Donat Wehner

## Abstract

The Medieval Ottonian dynasty had a lasting impact on European history. We obtained ancient genomic DNA from the purported remains of Otto I (912-973) and Heinrich (Henry) II (973-1024), the first and last emperors of this dynasty, preserved in the cathedrals of Magdeburg and Bamberg, respectively. Historical records attest that they were related as a great-uncle and a grandnephew via the paternal line. Whole-genome sequencing confirms such a relationship between the two individuals, as we identify a third-degree genetic relationship based on shared DNA segments and infer matching Y haplogroups. This genetic relatedness effectively identifies the remains of the two emperors. The authentication yields a valuable resource for refining and calibrating bio-archaeological methods. Because historical records provide the precise lifespans and dates of death of these individuals, their remains can serve as a “ground-truth” for methods such as radiocarbon dating and age-at-death estimates. They can provide calibration data to improve our understanding of the radiocarbon reservoir effects of Medieval elites. As the Ottonian lineage was closely linked to the mating networks of elites across Europe, the genomes of the two emperors are valuable resources for identifying other potential elite burials.

## Introduction

During the 10th and early 11th centuries, the Ottonian dynasty ruled the East Frankish Kingdom for more than 100 years (Althoff 2013). The dynasty had a profound influence on European history. The Ottonians assumed the title of Emperor and firmly associated it with the East Frankish Kingdom, thereby laying the foundation for what later became known as the “Holy Roman Empire” (Fried 2012). Because of their prominent role in European history, several of the Ottonian emperors, such as Otto I “the Great”, remain well known to this day. They also became projection surfaces, and the histories of Ottonian rulers have been frequently instrumentalized, including during the Nazi period in Germany (e.g., Kortüm 2014). In particular, Otto I’s father, Heinrich I, who was long anachronistically referred to as the first “German king”, was extensively instrumentalized by the National Socialists (Stahl 2013, Halle 2019). Interpretations of the research focus have changed over time - from national-state to national socialist appropriation to the consideration of contemporary circumstances without retrospective appropriation (Keller/Althoff 2008, 24–33).

This 10th-century Eastfrancian ruling family is known as the Liudolfingian, Ottonian, or Saxon dynasty. The “Liudolfingian” refers to Count Liudolf (died in 866), the “Ottonian” to his grandson Count Otto the Illustrious (died in 912), and “Saxon” to the medieval “Saxony” region. In the High Middle Ages, ‘Saxony’ encompassed the regions now known as Lower Saxony, Friesland, Saxony-Anhalt, and parts of Thuringia, whereas the present-day state of Saxony lay far outside medieval Saxony. Initially, the Ottonian family was of Saxon nobility (Becher 1996), and their ancestors were likely members of the Frankish functional elite under Emperor Charlemagne and his successors. The Ottonian family’s central base of power was the area around the Harz Mountains, known as the ‘Königslandschaft’ (King’s Landscape) (Schulze 1991).

The first Ottonian king (919) was Heinrich I (c. 876-936). He and his wife Queen Mathilde (c.896-968) had five children, one of whom was Otto I “the Great” (912-973). Otto I not only succeeded his father as king (936) but was also crowned emperor by the pope in Rome in 962. Earlier, the Frankish king Charlemagne had first claimed the Roman Empire for the Franks and proclaimed himself Emperor in 800, thus entering into competition with Byzantine rulers (Brühl 1995). Yet the last Western king carrying the title of “Emperor”, Berengar I. of Italy, had died in 924, before Otto I assumed this rank. Otto’s son (Otto II) and grandson (Otto III) subsequently also held the title of emperor. After Otto III died childless in 1002, he was followed by Heinrich II (973-1024), a grandnephew of Otto I, who became the fourth and last emperor of the Ottonian dynasty (1014). After Heinrich died childless as well (1024), the Salian dynasty took power through the descendants of a daughter from Otto’s first marriage to Queen Edgitha (c.910-946).

The burial sites of Ottonic emperors are frequently visited pilgrimage destinations and tourist attractions, and consequently targets of preservation efforts. However, the authenticity of the human remains remains unproven due to the scarcity of historical records from the millennium since the original burial and the potentially complex history of the osteological material. Numerous examples are known in which the graves of saints, bishops, and rulers were later opened and placed together in collective graves, resulting in the mortal remains being heavily intermingled (Meier 2002).

Our study focuses on the putative remains of Otto I and Heinrich II, the first and last Ottonian emperors, who were related as a great-uncle and a grandnephew (**Figure 1**) according to historical records (Glocker 1989, Hlawitschka 2006). That more than 1000 years have passed since their deaths, the disturbance of Otto’s grave (see **Text Box 1**) and the storage of Heinrich’s skull outside the grave (see **Text Box 2**) cast doubt on the authenticity of the remains. Exceptional circumstances in 2025 enabled us to collect samples from both individuals. To generate new evidence, we conducted an ancient DNA study of remains attributed to the two Ottonian emperors to determine whether the historically attested genealogy aligns with inferred genetic relatedness.

**Figure 1:**
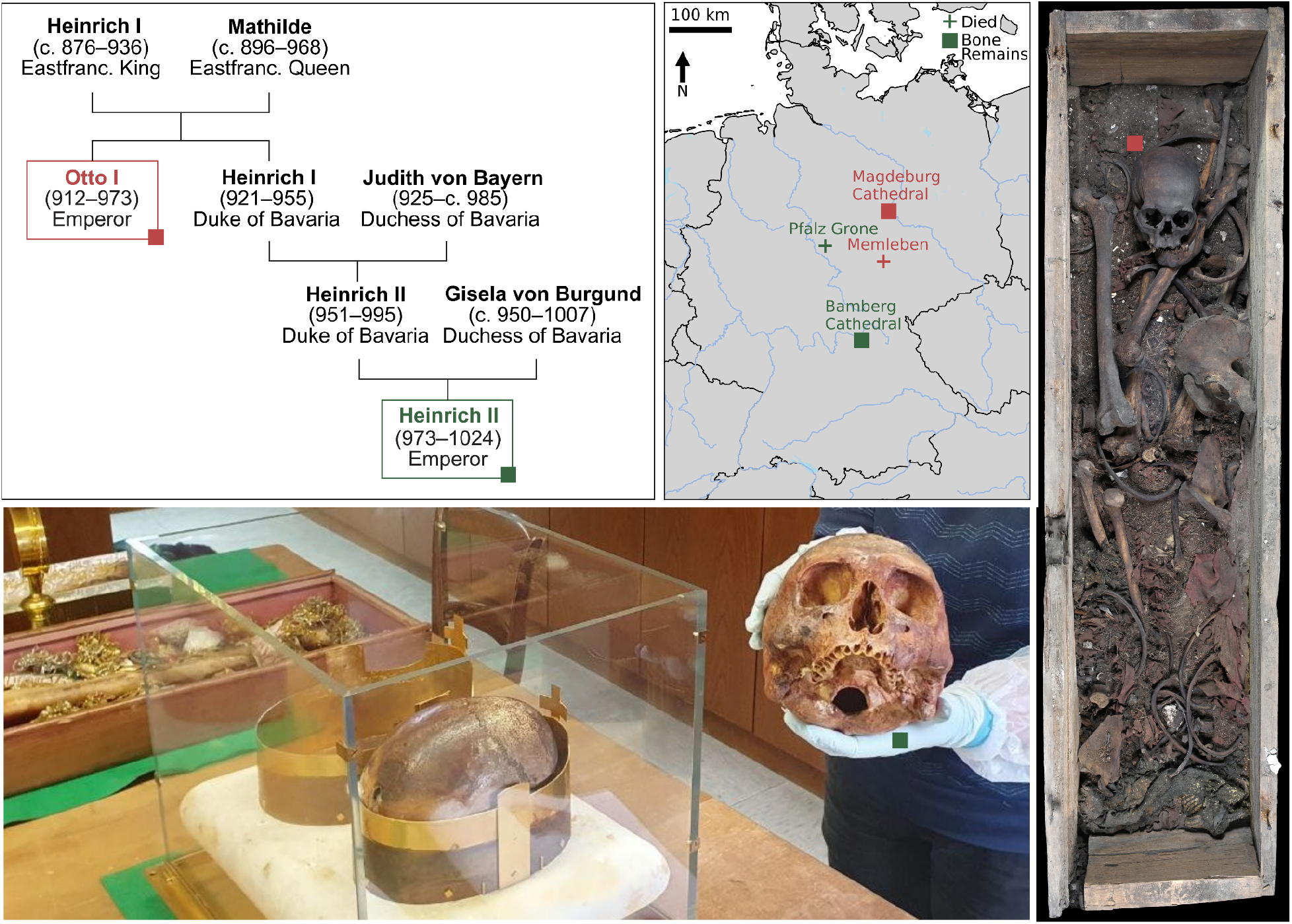
Historical Genealogy and sampled remains associated with Otto I and Heinrich II. The genealogy illustrates the third-degree kinship between Emperor Otto I and Emperor Heinrich II, as attested by medieval sources (Glocker 1989). The map depicts the burial locations of the bone remains associated with these two emperors (squares) and their places of death (crosses). The two photos depict the remains we sampled in 2025. Right Photo: remains in the grave of Otto I; Bottom Photo: remains associated with Heinrich II, a femur (in the box) and a skull (in the hands) together with the skull associated with his wife Kunigunde (in the glass box). The two photos are not covered by the Creative Commons license (for photo Otto I, © LDA Sachsen-Anhalt, Andrea Hörentrup).

### Emperor Otto I (912-973)

**Biography:** Otto was born on 23 November 912, most probably in Wallhausen (Freund 2013). During his lifetime, Otto participated in at least 50 sieges and 16 battles (Bachrach 2012). His epithet ‘the Great’ refers less to his physical size than to his distinction from his son Otto II, whom he had made co-king in 961 (Wozniak 2014). During the Battle of Lechfeld in 955, facing a Hungarian army, Otto vowed to establish an archbishopric in Magdeburg if he were victorious. He implemented this plan in 968 (Laudage 2001, Becher 2012).

**Remains:** He chose the Magdeburg Cathedral as the burial place for his first wife, Edgitha, and himself. After Otto died in Memleben on 7 May 973, most of his remains were brought to Magdeburg (his intestines stayed in Memleben, Schmitz-Esser 2023). Otto is buried in Magdeburg Cathedral, which he built but which has been rebuilt several times, including a complete reconstruction in the 13th century. His grave had last been officially opened in 1844. As documentation standards were still very low at that time, only a rough sketch of the tomb interior and a drawing of his skull were known from this opening. Inadequate security measures at Otto’s grave in the 19th century meant that there was a risk of long-term damage. Due to necessary renovation work, it was reopened in the summer of 2025, allowing an examination of his remains. The remains of Edgitha also remained in Magdeburg Cathedral and were examined in detail between 2008 and 2010. Due to poor preservation, no aDNA could be obtained from Edgitha (Meller et al. 2012).

**Text Box 1: Otto I**

### Emperor Heinrich (Henry) II (973-1024)

**Biography:** Otto’s brother, Heinrich (921-955), Duke of Bavaria, married Judith of Bavaria, and their marriage produced a son, Heinrich (951-995), who also became Duke of Bavaria. He was married to Gisela of Burgundy, and their son was Emperor Heinrich II (see **Fig. 1**). Heinrich was born on 6 May 973, probably in Bamberg (Weinfurter 2002). After the death of his great-cousin Otto III, he established himself as the new king in 1002. His marriage to Queen Cunigunde of Luxembourg can be described as a love match, as evidenced by the remarkable wording in various royal documents. Nevertheless, the couple remained childless, a fact contemporaries attributed to Heinrich (Weinfurter 2002). With the deaths of Heinrich in 1024 and his brother, Bishop Brun of Augsburg, in 1029, the male line of the Ottonian dynasty died out. Heinrich’s infertility is said to have been caused by the colic from which he apparently suffered. According to other sources, accidents were its cause. In 1024, his pain was so severe that he had to take several long breaks of weeks and months during a trip to the Harz region. He finally died on 13 July 1024 in the royal castle of Grone.

**Remains:** Heinrich’s remains were taken to Bamberg, where they were buried in the cathedral, which Heinrich had founded, as well as the whole diocese. Most of his body was relocated in the 16th Century into a famous tomb sculpted by Tilman Riemenschneider between 1499 and 1513. Yet his skull and some bones were kept outside because they were venerated as holy relics, as Heinrich had been canonized as a Catholic saint in 1146. The skulls of Heinrich and his wife Kunigunde, who was canonised in 1200 as well, were placed in metal reliquaries and regularly displayed during processions. At least one femur was transported to Rome at a later date. On the occasion of the femur’s repatriation to Bamberg in 2024, the opportunity opened up to take aDNA samples from Heinrich’s skull and femur at the beginning of 2025.

**Text Box 2: Heinrich II**

## Results

### Ancient DNA data generation

We sampled a Petrous Bone from the remains of Heinrich II in the Cathedral of Bamberg and an Incus Bone from the remains of Otto I in the Cathedral of Magdeburg. Starting from lysates of these samples, we extracted single-stranded ancient DNA libraries using a UDG half-treatment (see **Methods**). Whole-genome sequencing of both libraries and bioinformatic processing with a standard pipeline designed for aDNA (see **Methods**) indicated excellent DNA preservation, with high levels of human DNA carrying aDNA typical damage patterns (see **Tab. 1, Table S1, Fig. S1**). Deeper Sequencing of the two genomes resulted in an average genomic coverage of 0.996x and 0.576x, respectively. We inferred that both genomes are genetically male, based on the relative number of reads mapping to the Y and X chromosomes compared to the autosomes (see **Methods, Tab. 1**). We detected no significant ancient DNA contamination (see **Tab. 1**), with contamination estimates for the male X chromosome well below 2% (see **Methods, Tab. 1**).

**Table 1:** Summary statistics of the two ancient genomes. Endogenous DNA refers to the fraction of sequencing reads that map to the human genome after processing (including deduplication and filtering for read length and mapping quality). The imputation quality denotes the fraction of the so-called 1240k SNPs on chromosome 3 with maximum posterior genotype probabilities >0.99. The column “Avg. Genomic Coverage” is calculated as mean depth on the 1240k SNP set. The two columns “Rate X” and “Rate Y” denote the relative coverage of mapped reads on the X and *Y* chromosomes relative to the coverage on the autosome.

| Sampled Individual | Library Code | #Reads [Million] | Endogenous DNA | Avg. Genomic Coverage | Imputation Quality | Rate X | Rate Y | mtDNA Haplogroup | Y Haplogroup |
| --- | --- | --- | --- | --- | --- | --- | --- | --- | --- |
| Heinrich II | BMG001 | 672.46 | 10.28% | 0.996x | 0.941 | 0.5255 | 0.4878 | H7b2a | R1b-FTA63331 |
| Otto I | WSQ003 | 162.44 | 33.17% | 0.576x | 0.907 | 0.5205 | 0.4582 | K1a4a1a2b | R1b-FTA63331 |

### Genetic relatedness on the biological level of the 3rd degree

The whole-genome sequencing data enabled us to infer IBD segments (identity by descent), the stretches of DNA shared by two individuals and co-inherited from common recent ancestors. For this analysis, we first imputed diploid genotype probabilities using *GLIMPSE (Rubinacci et al. 2021)* and the modern 1000G reference panel (The 1000 Genomes Project Consortium 2015), and then used these data to infer long shared IBD segments with *ancIBD (Ringbauer et al. 2024)* (for a detailed description, see **Methods**). We detected several long IBD segments shared between the two genomes (**Fig. 2a-c, Table S1**). When comparing empirically observed IBD segment sharing with simulated IBD segment sharing generated using *Ped-sim* (Caballero 2019) across a large set of replicates of various relatedness clusters, we observe that the empirically observed IBD sharing clusters with simulated third-degree relatives, which are connected via one full-sibling pair (**Fig. 3**). Throughout, we denote degrees in the genetic sense, where each full sibling and parent-offspring pair adds one degree.

**Figure 2:**
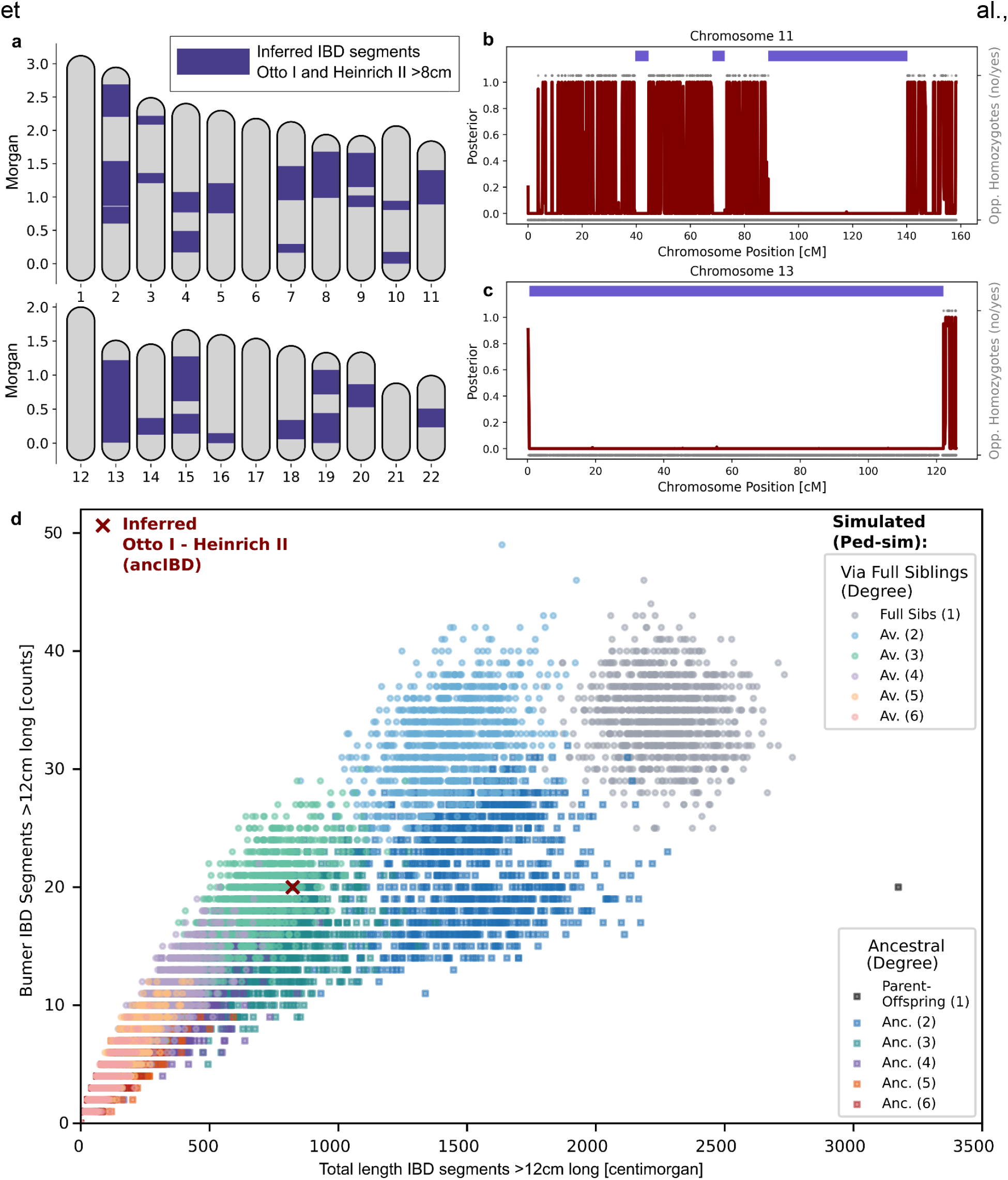
Shared IBD Segments between the Genomes of Otto I and Heinrich II. **a**: Karyotype depicting the positions of all inferred IBD segments >8cm long on all 22 autosomes. **b-c**: Posterior probability of not being in an IBD state (maroon) and inferred IBD segments >4cm long (depicted in blue) on Chromosomes 11 and 13, respectively. **d:** Comparison of empirical IBD sharing to simulated relationship classes. We simulated 1000 replicates of IBD segment sharing with *Ped-sIM* for various classes of relative pairs (see **Methods**). For each individual, we plot the sum (on the x-axis) and the number (y-axis) of inferred IBD segments.

**Figure 3:**
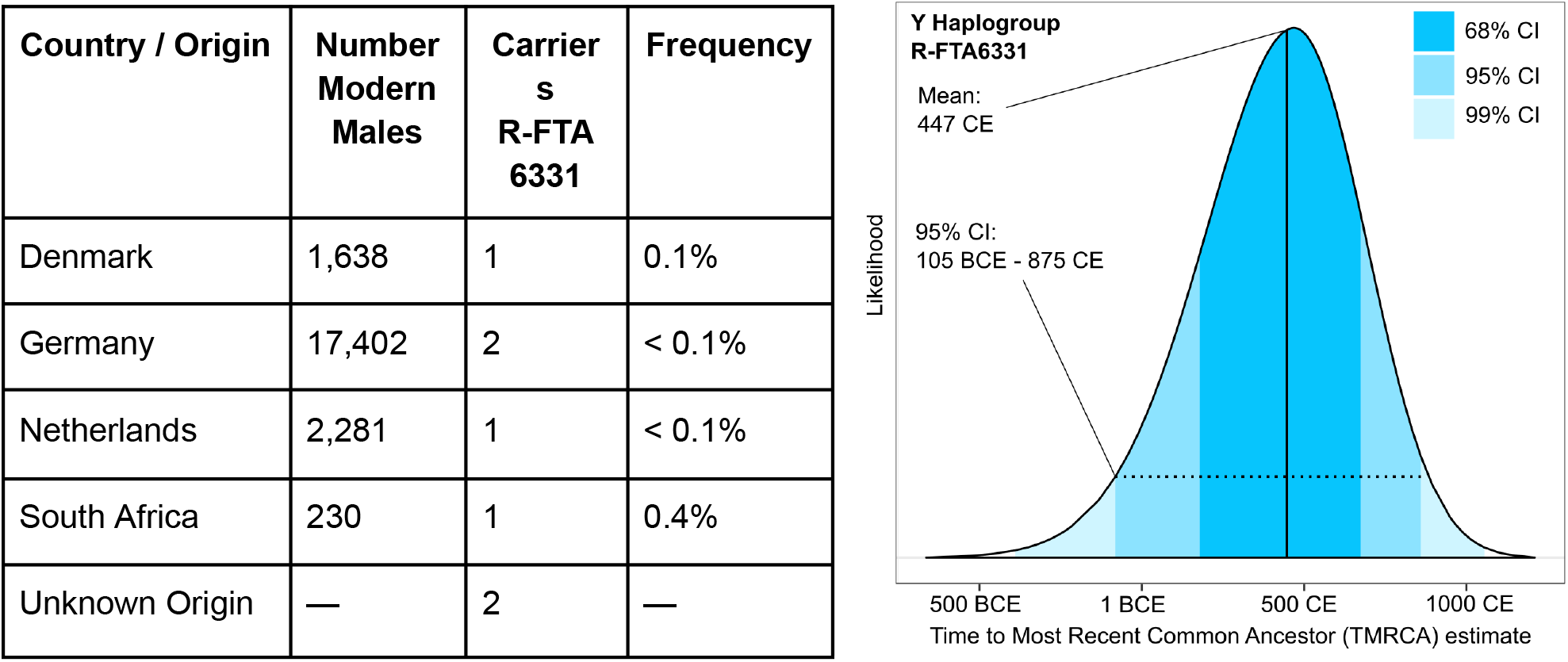
Overview of Y Haplogroup R-FTA63331. The Y Haplogroups of both the remains Otto I and Heinrich II can be confidently assigned to this Y haplogroup (see **Table S1E** for detailed SNP status). **Table**: We depict the frequency of this haplogroup in modern individuals in the FamilyTreeDNA Y-DNA Haplotree (version 2026.02.15). **Figure**: The depicted FamilyTreeDNA Time to Most Recent Common Ancestor (TMRCA) estimate is calculated based on SNP and STR test results from present-day DNA testers. The uncertainty in the molecular clock is represented in this probability plot, which shows the most likely time the common ancestor lived (adapted from familytreedna.com, March 2026).

### Uniparental Haplogroups

The genomic coverage of the Y chromosome (0.358x and 0.224x) and mitochondrial DNA (87.1x and 30.2x) enabled us to infer uniparental haplogroups for Heinrich II and Otto I, respectively. For these analyses, we queried the extensive *FamilyTreeDNA* database for present-day uniparental haplogroups and their associated derived markers (see **Methods**). Notably, this analysis revealed that both genomes share the same Y haplogroup, designated R1b-FTA63331 based on a derived SNP within this haplogroup (see **Table S1e**). This haplogroup is very rare in modern males, with only a single male carrier in the Netherlands, Denmark, Germany (at frequencies <0.1%), and South African males, and an inferred TMRCA (time to most recent common ancestor) of 448 CE (95% CI: 105 BCE-876 CE).

We inferred that the two individuals have different mitochondrial DNA (mtDNA) haplogroups (see **Table S1f** and **Table S1g**). Otto I is a carrier of K1a4a1a2b, a haplogroup occurring at low frequency across northwestern Europe (0.2-0.5%, see **Fig. 4a**). Heinrich II is a carrier of H7b2a, a haplogroup specific to modern central Europeans (peaking at 0.9% in Swiss and at 0.2-0.1% in French and Germans, see **Fig. 4b**).

**Figure 4:**
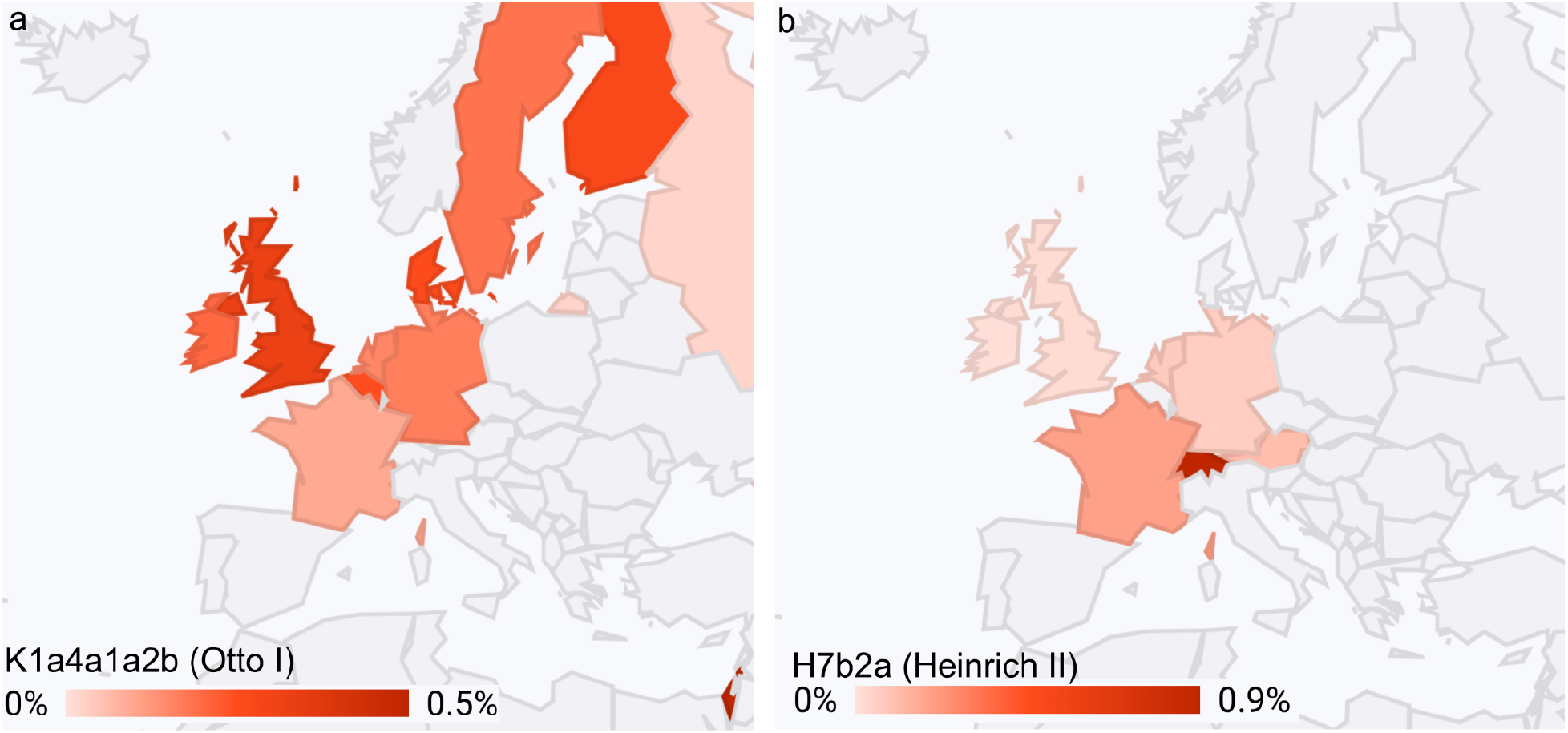
Geographic distribution of mitochondrial haplogroups. Geographic distribution of the two mtDNA haplogroups of the remains of Otto I (K1a4a1a2b) and Heinrich II (H7b2a) in the FamilyTreeDNA mtDNA database (adapted from familytreedna.com, March 2026).

## Discussion

The genomic data from the purported remains of Heinrich II and Otto I reveal a close biological relatedness between them. Notably, there are different approaches to labeling degrees of kinship in history, genetics, and jurisprudence. In the Roman *Codex Iustinianus* and in most modern legal systems, each procreation counts as one degree, whereas the canonical count follows the first common ancestor, as in the genetic approach used here. The number and size of shared IBD segments indicate that the relationship is (in the genetic sense) third-degree and that the line passes through intermediate full siblings. Moreover, the shared Y haplogroup provides evidence that the relationship traces back through an unbroken male lineage. Together, the patterns perfectly match their historically attested relationship, where Otto I is the great-uncle via the paternal line of Heinrich II.

These results effectively authenticate the remains of both emperors. It would be highly implausible that not only one but both of the two emperors’ remains were mixed up, and that, moreover, the replacements were themselves a pair of third-degree relatives via the paternal line. We can therefore, with probability bordering on certainty, conclude that both remains are indeed those of Otto I and Heinrich II. Moreover, our findings rule out any extrapaternal events in the lineage linking the last Ottonian emperor (Heinrich II) to the first (Otto I).

The authentication of these two human remains of Medieval elites is not only important in itself (e.g., for future preservation efforts) but also yields a valuable resource for refining and calibrating Bio-Archaeological methods. Because historical records provide the precise lifespan and date of death, the over 1000-year-old remains of Heinrich II and Otto I can serve as a “ground-truth” for methods such as radiocarbon dating and age-of-death estimates. For instance, the two known remains can provide important calibration data to improve our understanding of the radiocarbon reservoir effects of Medieval elites.

The high-quality genetic data we generated here will also be useful for studies of other Medieval elite contexts. As the Ottonian lineage was closely linked to the mating networks of Medieval nobility - such as the Capetians in France, the Burgundian kings, and the Salian dynasty - and had close ties throughout Europe, the genomes of the two Ottonian emperors can serve as valuable resources for identifying and authenticating potential Medieval elite burials.

## Supplementary Material

**Table S1**: Tables with information about ancient DNA data.

## Methods

### DNA Data Generation

The DNA extraction and library preparation were performed at the Department of Archaeogenetics (MPI-EVA) in Jena, Germany. Sequencing was performed in the Core Unit (CU) of MPI-EVA in Leipzig using an Illumina NovaSeqX Plus sequencing platform. DNA was extracted using a silica-based method optimized for the recovery of short DNA fragments (Dabney et al. 2013), following a modified version of a published protocol (https://doi.org/10.17504/protocols.io.baksicwe). First, bone powder samples were subjected to decalcification and denaturation by adding 1 mL of extraction lysis buffer (0.45 M EDTA, pH 8.0, 0.25 mg/mL proteinase K, 0.05% Tween-20). Then, 125 μl were aliquoted from the resulting lysate and purified using an automated handling system (Bravo NGS Workstation B, Agilent Technologies) with silica-coated magnetic beads and binding buffer D, as previously described (Rohland et al. 2018). The final elution volume was 30 μl, and the entire product was used for library preparation. The remaining lysate was frozen (-25 °C) and re-aliquoted as described if more than one DNA library was requested per sample. An extraction blank was included with each extraction plate. An automated version of the single-stranded DNA library preparation (Gansauge et al. 2017) was applied to all DNA samples, following the published protocol (Gansauge et al. 2020). The library preparation plate included an extraction blank and additional negative controls (library blanks). Library yields and the efficiency of library preparation were determined using a single quantitative PCR assay (Gansauge et al.(Gansauge et al. 2020). The libraries were amplified to a plateau by performing 35 PCR cycles and tagged with pairs of library-specific indices using AccuPrime Pfx DNA polymerase ((Gansauge et al. 2020). Amplified libraries were purified using SPRI technology (DeAngelis et al. 1995) as described in (Gansauge et al. 2020).

### Bioinformatic Processing and Quality Control

We processed demultiplexed sequencing data using *leeHom* [v1.1.5] (Renaud et al. 2014) to merge paired-end reads and trim Illumina adaptors. Forward, reverse, and merged reads were mapped to the human reference hs37d5 using the Burrows–Wheeler aligner (BWA; v.0.7.12) (Li and Durbin 2009) with parameters -n 0.01 -o 2 -l 16500 (seeding disabled). We processed the output files in Binary Alignment Map (BAM) format using the *nf-core/eager* Nextflow pipeline [v2.4.6] (Fellows Yates et al. 2021; Ewels et al. 2020) using a custom configuration file from MPI-EVA (https://github.com/nf-core/configs/blob/master/docs/eva.md) and with parameters set as listed in **Table S1c**.

This pipeline outputs processed.bam files with duplicate reads removed and quality-filtered reads (base quality > 30, mapping quality > 25, and read length > 30). These processed.bam files served as the basis for our summary statistics (**Tab. 1**) and for subsequent IBD analysis.

To quantify human DNA contamination, we applied two methods to BAM files filtered for mapping quality >30: *hapCon (Huang and Ringbauer 2022)* and *Schmutzi (Renaud et al. 2015)*. Using a haplotype-copying imputation approach, *hapCon* estimates contamination in aDNA data from male individuals with a lower coverage threshold and lower uncertainty than similar tools for male contamination estimation. *Schmutzi* estimates sample contamination in mitochondrial reads (Furtwängler et al. 2018).

### Imputation and detecting shared haplotypes

To infer pairwise shared identity-by-descent (IBD) segments, we used ancIBD (Ringbauer et al. 2024), software designed to detect such shared haplotypes in aDNA data. We began with processed BAM files containing all aligned reads and then imputed genotype probabilities using *GLIMPSE (Rubinacci et al. 2021)* with the 1000 Genomes reference panel, following a previously described imputation pipeline (Ringbauer et al. 2024). Using the recommended default settings and cutoffs for *ancIBD*, we inferred all IBD segments longer than 8 cM. The imputation quality for both genomes is substantially above the recommended ancIBD quality cutoff of 0.7 (see **Tab. 1**).

### Simulating IBD segment sharing

We simulated IBD segments between relatives using the software *Ped-sim* (v1.0.6) (Caballero et al. 2019), building on the approach described by (Ringbauer et al. 2024). For each degree of relatedness up to the sixth, we simulated 1000 pairs of individuals using the sex-specific genomic map from (Bhérer et al. 2017) and a recombination interference model in *Ped-sim*. To compare those to empirical IBD inferred with ancIBD, we applied the same IBD segment filter that we applied to post-process *ancIBD* output: We merged adjacent IBD segments, such as occurring when there is a switch between IBD1 and IBD2 states that are output as separate segments in *Ped-sim*, into a single contiguous one and removed IBD segments that have a density of 1240k SNP less than 220 per centimorgan. We note that the latter filter removes chromosomes 19 and 22 if they are completely IBD (e.g., in parent-offspring).

For each degree of relationship, we simulated two types of relatives: ancestral relationships (such as parents and grandparents) and relationships via full sibs (such as full sibs or uncles/aunts). These two relationship types have different distributions of IBD lengths at the same degree of relatedness because, for the same degree of relatedness, the number of meioses in relationships via full sibs increases by one, whereas the number of shared haplotype ancestors increases from two to four.

### Uniparental analysis

We realigned Y chromosome-mapped reads to GRCh38 using *BWA-ALN* (v0.7.17) for comparison with the FamilyTreeDNA Y-DNA Haplotree (version 2026.02.15), which has 100,286 branches. Duplicates were marked using *GATK* (v4.2.2.0) *MarkDuplicates*. Variant calling was performed using *BCFtools* (v1.22) in gVCF mode, requiring base and mapping qualities ≥30. We compared variants and non-variant sites to the phylogenetic tree, and each candidate node was scored based on the number of positive and negative SNP calls leading to it. All covered variants in the non-recombining region of the Y chromosome (NRY) were considered, and we manually inspected aligned reads for all diagnostic SNPs to confirm their positive, negative, or indeterminate status.

We realigned mtDNA-mapped reads to *RSRS* using *BWA-MEM*. Duplicates were marked, and variants were called in the same way as for Y-DNA. Variant calls with low-level heterozygous calls for expected deamination artifacts (C>T and G>A variants with the T/A allele depth below 20%) were adjusted to prevent them from being called as false-positive heteroplasmies. Mitochondrial base pairs without any coverage or with a genotype quality below 13 were treated as no-calls. The resulting sequence was compared against the FamilyTreeDNA Mitotree (version 2025.12.13), which has 53,856 branches, using *Haplogrep* (v3.2.1).

## Code Availability

The code used to analyze and visualize the results presented in this work is publicly available on GitHub (https://github.com/hringbauer/medieval_elites.git).

## Data Availability

The newly generated genomic data will be made publicly available at publication in a Scientific journal. Until then, raw and processed genomic data are available upon request for replication or to identify other Medieval elites.

## Declaration of interests

We declare no competing interests.

### Author Contributions

Our annotation of author contributions follows the CRediT Taxonomy labels (https://credit.niso.org/). When multiple authors fulfil the same role, we identify their degree of contribution as either lead, equal, or support.

Conceptualisation - TW, JF, JK, HR, JK, HM, DW Data curation - RAB

Formal analysis - HR

Funding acquisition - JK, HM, HR Investigation - RAB, HR Visualisation - HR, TW

Writing (original draft) - HR, JF, TW

Writing (review & editing) - input from all co-authors

## Acknowledgements

We would like to thank the Archbishop of Bamberg, Herwig Gössl, and the entire cathedral chapter for the opportunity to sample ancient DNA from Emperor Heinrich II, as well as Dr. Birgit Kastner for her helpful support. We would like to express our gratitude for the excellent cooperation with the Saxony-Anhalt Cultural Foundation, the Magdeburg Cathedral Parish, the Protestant Church in Central Germany, and the state of Saxony-Anhalt.

HR and JK were supported by funding from the Max Planck Society and the Max Planck-Harvard Research Center for the Archaeoscience of the Ancient Mediterranean (MHAAM).

## Supplementary Figures

**Figure S1:**
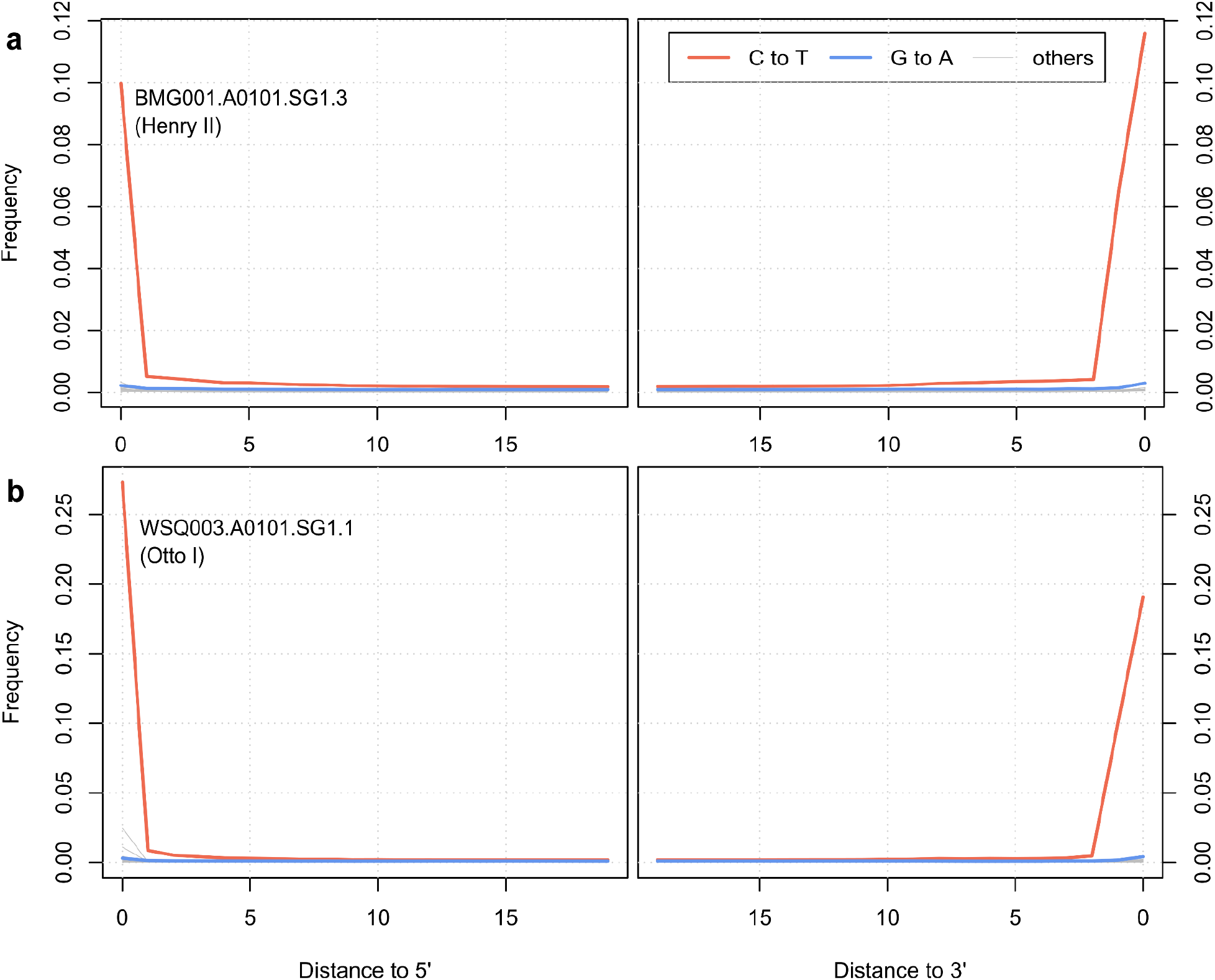
Ancient DNA damage patterns. We depict the ancient DNA damage patterns for the Heinrich II DNA library (panel **a**, top) and the Otto DNA library (panel **b**, bottom). We plotted the damage patterns calculated from all human DNA fragments longer than 35 base pairs. The damage pattern is typical for UDG-half-treated libraries, with C->T damage appearing in the 2-3 bases at the end of sequences.

## Notes

### Competing Interest Statement

The authors have declared no competing interest.

